# Study of Glucometer Precision and Antimalarial Therapy Interference on Glucose Readings in Nigeria

**DOI:** 10.1101/2023.05.28.542634

**Authors:** Ezinwanne Jane Ugochukwu, Adaobi Uchenna Mosanaya, Victor Nnanna Nweze, Marydith Ifeoma Chukwu, Chukwuemeka Michael Ubaka, Mohammed Alawami

## Abstract

Malaria-infected diabetic (MID) patients have become one of the highest diabetic population with comorbidities in Nigeria. While the artemisinin-based combination therapy (ACT) are used for malaria treatment, glucometers are used to monitor their blood glucose levels. However, some drugs are known to interfere with the accuracy of glucose readings leading to misdiagnosis. In this study, we investigated the interferences of artemether and lumefantrine as well as other known drug interferents on the accuracy of glucose readings in three different glucometers commonly used in Nigeria **(Accu-Answer®, VivaChek® and GlucoDr®)**. Blood sample was collected from a healthy volunteer, aliquots of the blood with pure samples of artemether and lumefantrinepowders as well as other known drug interferents were prepared in normal and low pools. The precision study on the glucometers were determined. We find that Accu-Answer**®** was the only precise glucometer considering its coefficient of variation. The influence of the artemether and lumefantrine was mild in two glucometers andthe predicted binding affinities showed possible binding interactions that may have caused their interferences.

## 1. Introduction

Diabetes Mellitus (DM) is a metabolic disorder characterized by abnormal hyperglycemia resulting from an anomaly in glucose metabolism due to insulin deficiency and dysfunction of organs [1]. Over the years, DM has shown a significant increase in its demographic prevalence worldwide and as a result, there is a rapid increase in reported prevalence, which has become a socioeconomic and health challenge in manycountries [2].

In 2018, the World Health Organization (WHO) reported about 1.5 million deaths due to DM. As of 2021, the International Diabetes Federation (IDF) reported that 1 in 22 adults in Africa have diabetes leading to an alarming population of 24 million with a death toll of 416,000. The diabetic populace is predicted to increase to 55 million (129%) by 2045 [3]. Nigeria, being the largest country in Africa, ranks second among the top 5 countries in Africa with the highest reported diabetic cases of 3.6 million (3.7% of adult population)in 2021 for ages between 20 – 79 years [3]. However, glycemic control can be achieved by following lifestyle monitoring (strict healthy diet, regular exercise, and blood glucose (BG) monitoring) and medication as prescribed [4,5,6].

Using a Point-of-Care (POC), glucose monitoring is critical to ensure that the glycemic index of patients is within the acceptable threshold. However, glycemic index regularly present differently to diverse range of demography and clinical situations [7]. Most importantly, the accuracy of glucometers are being compromised by interferences influenced by regularly consumed medicines. Several studies have reported that acetaminophen (paracetamol) is a known glucometer interference that spikes BG readings beyond the actual BG level [7,8]. This is because its phenolic moiety is easily oxidized to yield more hydrogen peroxides that issensed by the glucometer and provides false readings in glucometers [9]. Other known interferences are ascorbic acid (vitamin C) and maltose can lead to false increase in glucometer results or errors in glucose measurements using the glucometer. As a result of these errors in glucometer, healthcare workers could present misdiagnosis of the patients [10], thereby causing life-threatening prescriptions for the patients.

Consequently, the increase in malaria cases in sub-Saharan Africa, especially in Nigeria, is of great concern for global health. According to the WHO World malaria report 2021, Nigeria accounted for the highest reported case (27%) which contributed to 96% of global malaria cases [11]. The combined increase in malaria and diabetes in Nigeria constitutes a major public health burden, especially with the high consumption of variousmedications for malaria-infected diabetic patients. To address this malaria catastrophe, the WHO adopted the Global Technical Strategy for Malaria 2016 – 2030 which was updated in 2021 [11]. The strategy stated that artemisinin -based combination (ACTs) therapy such as artemether-lumefantrine (A-L) remains the first-line of treatmentfor uncomplicated malaria because it is highly effective. Several studies reported symptomatic and asymptomatic cases of malaria-infected individuals with diabetes under antimalarial medication in Nigeria [12-16]. Considering the high intake of antimalarial drugs by diabetic patients in Nigeria, there is no study that has shown the possible interference of ACT drugs in glucometer measurement.

Therefore, this study aims to show the infleunce of a commonly used antimalarial ACT on three glucometers used in Nigeria based on glucose oxidase (GOx), glucose dehydrogenase (GDH)-flavin adenine dinucleotide (FAD), and GDH-pyrroloquinoline quinone (PQQ) assays.

## 2. Materials and methods

### 2.1 Materials

**Pure Drug Samples;** Dextrose powder, Maltose powder, Ascorbic Acid powder, Acetaminophen Powder, Artemether Powder, Lumenfantrine Powder. **Reagent/ Blood Sample;** Distilled water and Blood samples. **Glucometers and Test Strips;** Accu Answer^®^ plus 200 teststrips, Viva Chek^®^ plus 200 test strips, Gluco Dr^®^ plus 200 test strips. **Analytical Software;** Microsoft Excel version 16. **Laboratory Equipments;** Test tubes, Beakers, Stirrer, Glass slide, Micropipette, weighing balance, Measuring Cylinder and Lithium Heparin tubes.

### 2.2 Methods

#### 2.2.1 Ethical approval

The ethical approval for this study was granted by the Research Ethics Committee of the Faculty of Pharmaceutical Sciences, University of Nigerian, Nsukka, Enugu state, Nigeria. The confidentiality of the volunteer was ensured throughout the study.

#### 2.2.2 Blood collection

The informed consent form was filled by the healthy volunteer and the volunteer was screened for any disease condition mainly Human Immunodeficiency Virus, Hepatitis B&C, Diabetes, syphilis or any metabolic disorder. After the blood screening, 120ml of blood was collected from the healthy volunteer directly into 30 lithium heparin tubes.

#### 2.2.3 Glucometer Selection

A survey of glucometers commercially available and mostly used in Nigeria was conducted at three (3) pharmacies in Enugu state. Three glucometers were selected based on the frequency of its use by clients, accessibility of glucometer strips and the cost. The selected glucometers were Accu Answer^®^, VivaChek^®^ and GlucoDr^®^

#### 2.2.4 Study Procedure

##### 2.2.4.1 Precision study of Glucose

The precision study was carried out with dextrose. Ninety-Nine (99µl) of the whole blood of both normal and low pool were spiked with 1µl of various concentrations of dextrose (20, 40, 60, 120, 400 mg/dl) that had been initially prepared by serial dilution. Each spiked sampled, together with the baseline samples from the normal and low pools were measured on three different brands of enzyme-based glucometers; Accu -Answer^®^ containing GDH-PPQ, GlucoDr^®^ containing GDH-FAD, and Vivachek^®^ containing Glucose oxidase. The above was repeated three times to obtain the mean value. The same step was repeated on the following day.

##### 2.2.4.2 Interference Study

After obtaining ethical approval from the Research Ethics Committee of the faculty of pharmaceutical sciences university of Nigeria, Nsukka, 120ml of whole blood wasdonated from a healthy donor without diabetes or any metabolic disorder after filling the blood consent form and distributed directly into 30 lithium heparin tubes. After which, thebaseline readings were done using the three glucometers, Accu -Answer^®^ containing GDH-PPQ, GlucoDr^®^ containing GDH-FAD, and Vivachek^®^ containing GOx. Some of the lithium heparinized whole blood were left at room temperature for more than 6 hoursto allow blood glucose reduction from glycolysis to obtain the low blood glucose pool. Meanwhile, for the remaining lithium heparinized whole blood (normal pool), a number of aliquots were diluted based on the concentrations of the interference (Ascorbic acid, Maltose, Acetaminophen, Artemether and Lumefantrine). Using a micropipette, 1µl of each aliquot of interferents were added to 99 µl of whole blood sample, the glucometer strip was used to collect the blood through capillary action [7,17]. Three readings were obtained per concentration and this was repeated for other glucometers. This was also conducted for the low pool as shown in figure 1.

**Figure 1:**
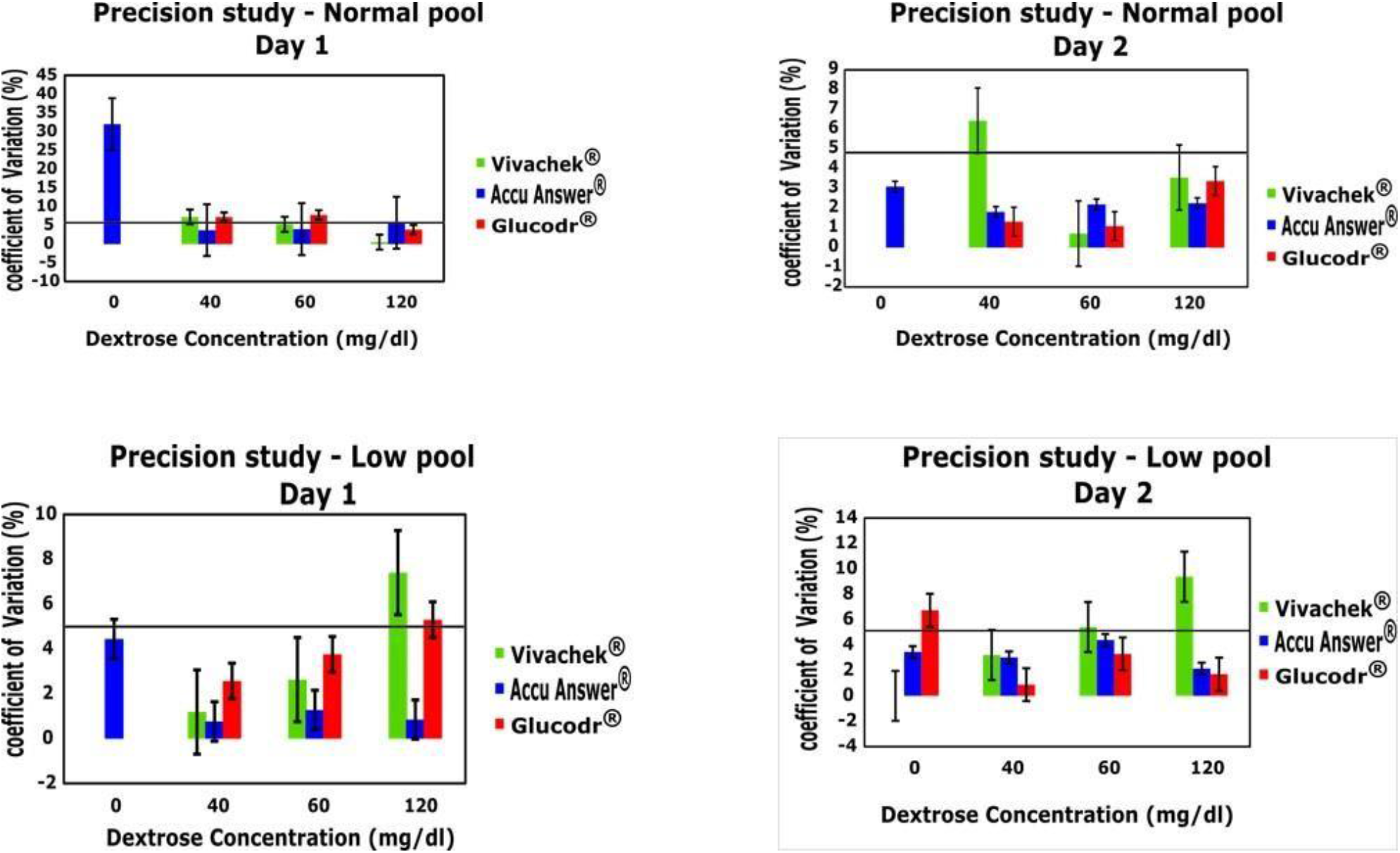
Precision study of three glucometers used in Nigeria. Coefficient of variation for precision study of the three glucometers – Vivachek^®^, Accu Answer^®^ andGlucodr^®^ – after two days. The Coefficient of variation for this precision evaluation is 5%.

##### 2.2.4.3 Molecular docking

###### Protein preparation

The crystal structure of Glucose oxidase (1CF3), Glucose dehydrogenase FAD (4YNU), Glucose dehydrogenase PQQ (1CQ1) was retrieved from Protein Data Bank (PDB) repository. The protein was prepared using the protein preparation wizard panel of Glide (Schrödinger Suite 2021-2) where bond orders were assigned, hydrogen added, disulfide bonds created, while missing side chains and loops were filled using prime. Water molecules beyond 3.0 Å of the heteroatoms were removed and the structure was minimized using OPLS2005 and optimized using PROPKA [18]. Subsequently, the receptor grid file was generated to define the binding pocket for the ligands.

###### Ligand preparation

Artemether and Lumefantrine were prepared for molecular docking using LipPrep module (Schrödinger Suite 2021-2). Low-energy 3D structures with correct chiralities were generated. The possible ionization states for each ligand structure were generated at a physiological pH of 7.2 ± 0.2. Stereoisomers of each ligand were computed by retaining specified chiralities while others were varied.

###### Receptor grid generation

Receptor grid generation allows defining the position and size of the protein’s active sitefor ligand docking. The scoring grid was defined based on the co-crystalized ligands (Flavin-adenine dinucleotide, D-glucano-1,5-lactone and pyrroloquinoline quinone) using the receptor grid generation tool of Schrödinger Maestro 12.8 the van der Waals (vdW) radius scaling factor of nonpolar receptor atoms were scaled at 1.0, with a partial chargecut off of 0.25.

###### Protein-Ligand Docking

Glide tool of Schrödinger Maestro 12.8 was used to perform the molecular docking studies using the generated receptor grid file. The prepared ligands were docked using standardprecision (SP), setting ligand sampling to flexible, with the ligand sampling set to none (refine only). The vdW radius scaling factor was scaled at 0.80 with a partial charge cut-off of 0.15 for ligand atoms.

### 2.3 Quantification, Statistical and Visualization Analysis

The mean values were calculated using Microsoft excel, other descriptive and inferential statistics were performed and graphs extrapolated. All error bars are reported as st33andard error 0of mean (±SEM). 2D view of the ligand-protein complexes was used to identify the chemical bonding, and interactions with amino acid residues in the binding poses.

## 3. Results and Discussion

### 3.1 Glucometer principle

This study evaluated the effects of artemether and lumenfantrine as interferents in glucometers such as Accu -Answer (GDH-PQQ), GlucoDoc (GlucoDr) (GDH-FAD) and Vivachek (GOx). The precision study was carried with dextrose (glucose). Other known interferences such acetaminophen, ascorbic acid and maltose were used to validate theprotocol for this study.

### 3.2 Precision study of three glucometers

Dextrose was used to assess the precision of three glucometers commonly used in Nigeria. From Figure 1, the coefficient of variation (CoV) was fixed at 5%. This means that for a glucometer to be considered precise, it must have a CoV below 5% across all concentrations for two days. Among the three glucometers, Accu-Answer^®^ was the only glucometer which retained CoVs within the threshold in both normal and low pool for twodays. This means that Accu-Answer^®^ glucometer is more precise in glucose readings than Glucodr^®^ and Vivachek^®^ glucometers. GDH-PQQ enzyme which is embedded in Accu -Answer is known to oxidize glucose and other sugar moieties [ 1 0 ]. This is consistent with Cho et al where Accuchek^®^ showed the high increase in glucose reading among two other glucometers [17]. Conversely, GOx enzyme in Vivachek^®^ favorably catalysis only glucose by binding to the five hydroxyl groups in the conserved binding pocket.

### 3.3 Interference study of the three glucometers

#### 3.3.1 Ascorbic acid interference

In Figure S2, all glucometers exhibited significant an increase in glucose readings at all concentrations of ascorbic acid. In normal pool, Vivachek^®^ and GlucoDr^®^ glucometers showed significant increase in the glucometer readings, with Vivachek^®^ expressing the highest glucose reading at 5 g/dL of ascorbic acid. In low pool, we observed significant increase in glucose readings for all glucometers with peak readings at 5 g/dl of low-pool ascorbic acid. Therefore, ascorbic acid interference was observed in all glucometers in both normal and low pools. This is because ascorbic acid is an antioxidant that can be oxidized to produce electrons and increase electrochemical signaling at the strip surface. Ascorbic acid contributes to oxygen depletion and hydrogen peroxide production when glucose is also catalyzed by glucose oxidase [19,20]. This observation is consistent with previous studies that reported ascorbic acid as a potential interferant [17,21,22].

#### 3.3.2 Acetaminophen interference

In Figure S2, various concentrations of acetaminophen were examined for the interference potential on the three glucometers for both normal and low pools. In the normal pool, Accu-Answer^®^ and Glucodr^®^ showed a significant increase in glucose readings with acetaminophen while we observed Vivachek^®^ showed an insignificant decrease in glucose readings. In low pool of acetaminophen, we observed increase in glucose readings in Accu-Answer^®^ and GlucoDr^®^, respectively while Vivachek^®^ exhibited no readings. Our findings do not support the observations of Vanavanan et al. (2010) where they reported that acetaminophen produced negligible interference in Accu -Chek^®^ (GDH assay) for low and high glucose levels while Optium^®^ (GDH assay) and Nova StatStrip^®^ (GOx assay) expressed significant negative and minor interferences, respectively [20]. However, our study supports several studies that proved that acetaminophen interfered with BG measurement [7,8,23,24]. This is because the phenoic moiety of acetaminophen is directly oxidized after diffusing the porous membrane of the electrodesurface, thus spiking the glucose readings [8,20] and leading to misdiagnosis of BG levels.

#### 3.3.3 Maltose Interference

Maltose is a disaccharide of two glucose moieties with an α-1,4-glycosidic bond. In Figure S3, we studied for the influence of maltose on the glucose readings of three glucometers. We observed significant increase in glucose readings with an increase in maltose concentrations in Accu-Answer^®^, GlucoDr^®^ and Vivachek^®^. In the normal pool, Accu-Answer^®^ and GlucoDr^®^ glucometers expressed higher glucose readings than Vivachek^®^. This shows that maltose exhibited mild interferences in these two glucometers. Our study is in contrast with the observations of Cho et al where they reported no significant interference by all maltose concentrations three different glucometers brands. However, these observations validate the claims that GDH reacts favorably with maltose [20,25] to interfere on the accuracy of glucose readings. Vivachek^®^ reported minor significant positive interference at normal concentrations. This may be asresult of the low glucose oxidase activity for maltose due to the free hydroxyl groups in maltose; four hydroxyl groups and the suppose fifth hydroxyl group at first carbon (C1) forming an α-1,4-glycosidic bond with another glucose moiety at the C4. The maltose interference maybe as a result of the binding of the free hydroxyl groups in the maltose.

#### 3.3.4 Artemether Interferences

Artemether is a commonly used antimalarial medicine through a combination therapy in Nigeria as recommended by the WHO. In this study (Figure 2), we investigated for the interference of artemether at 2, 4, 6 and 8 g/dL of artemether on the glucose readings of the three glucometers. Interestingly, all glucometers showed no significant increase in glucose readings in normal and low pools. Accu-Answer^®^ and Glucodr^®^ glucometers exhibited mild interferences in glucose readings while there were no significant interference in Vivachek^®^ glucometer. In low pool of artemether, there were no glucose readings in Vivachek^®^ glucometer. The specificity of the GOx in Vivachek^®^ glucometer may be responsible for the lack of artemether interference in normal and low pools. The increased artemether interferences in GDH assays may be due to the broad substrate specificity of GDH [26], which can yield false positive glucose readings.

**Figure 2:**
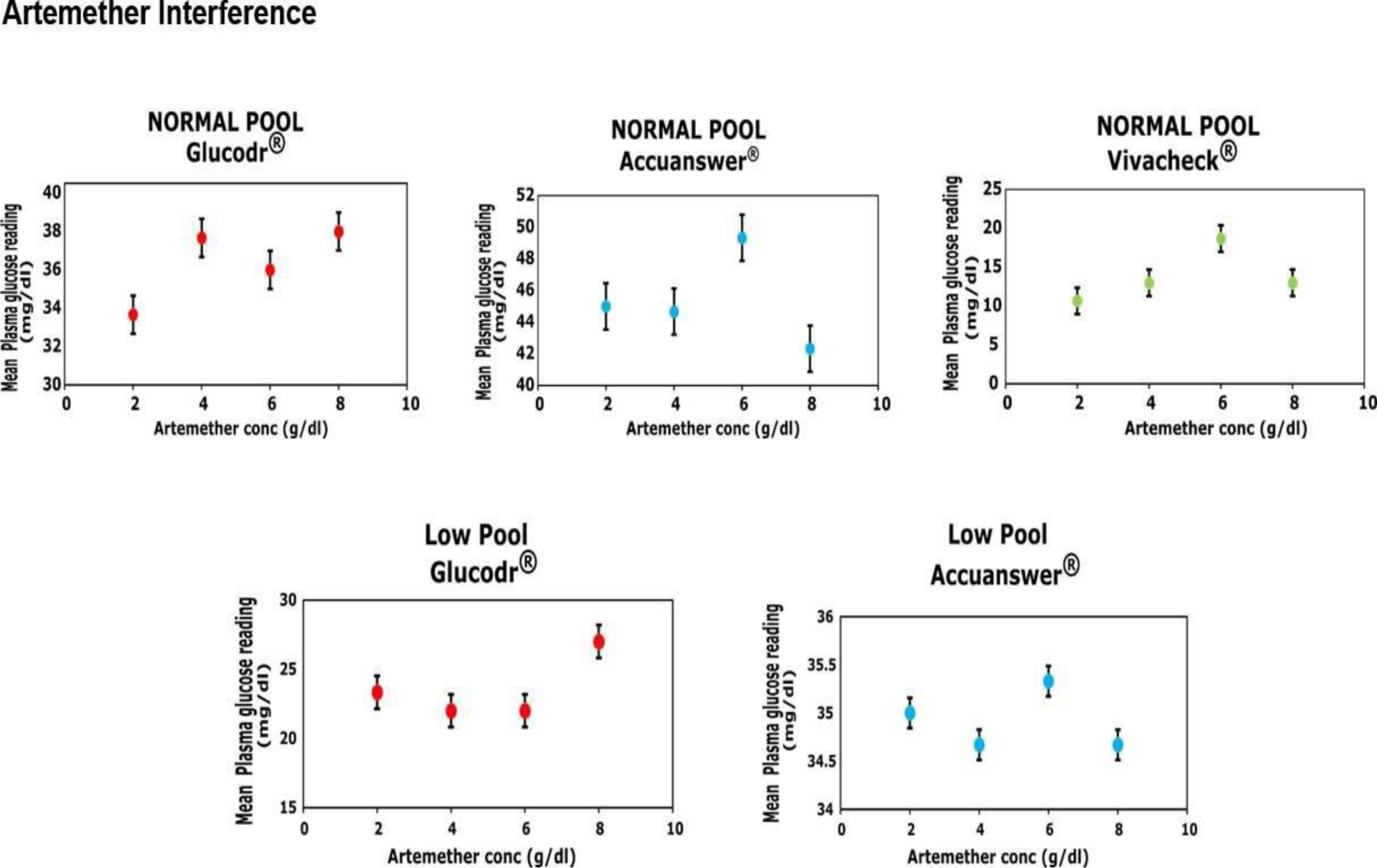
Influence of Artemether in glucose level measurement in three models of glucometers Vivachek^®^ Accu Answer^®^ and Glucodr^®^ – in normal and low pools.

#### 3.3.5 Lumefantrine interferences

Lumefantrine is an antimalarial drug that is widely used as a combination therapy in Nigeria according to the recommendation of the WHO. Figure 3 shows the outcomes of the influence of lumefantrine on glucometer readings in the three glucometers used in Nigeria. We observed that only Accu-Answer^®^ glucometer exhibited non-significant increase in glucose readings and may represent mild interference of lumefantrine in normal pool. We observed no significant increase in glucose readings in Vivachek^®^ glucometer but without any interference. In low pool of lumefantrine, mild interference of glucose was observed in Accu -Answer^®^ glucometer. However, there were no lumefantrine interference in Glucodr^®^ and Vivachek^®^.

**Figure 3:**
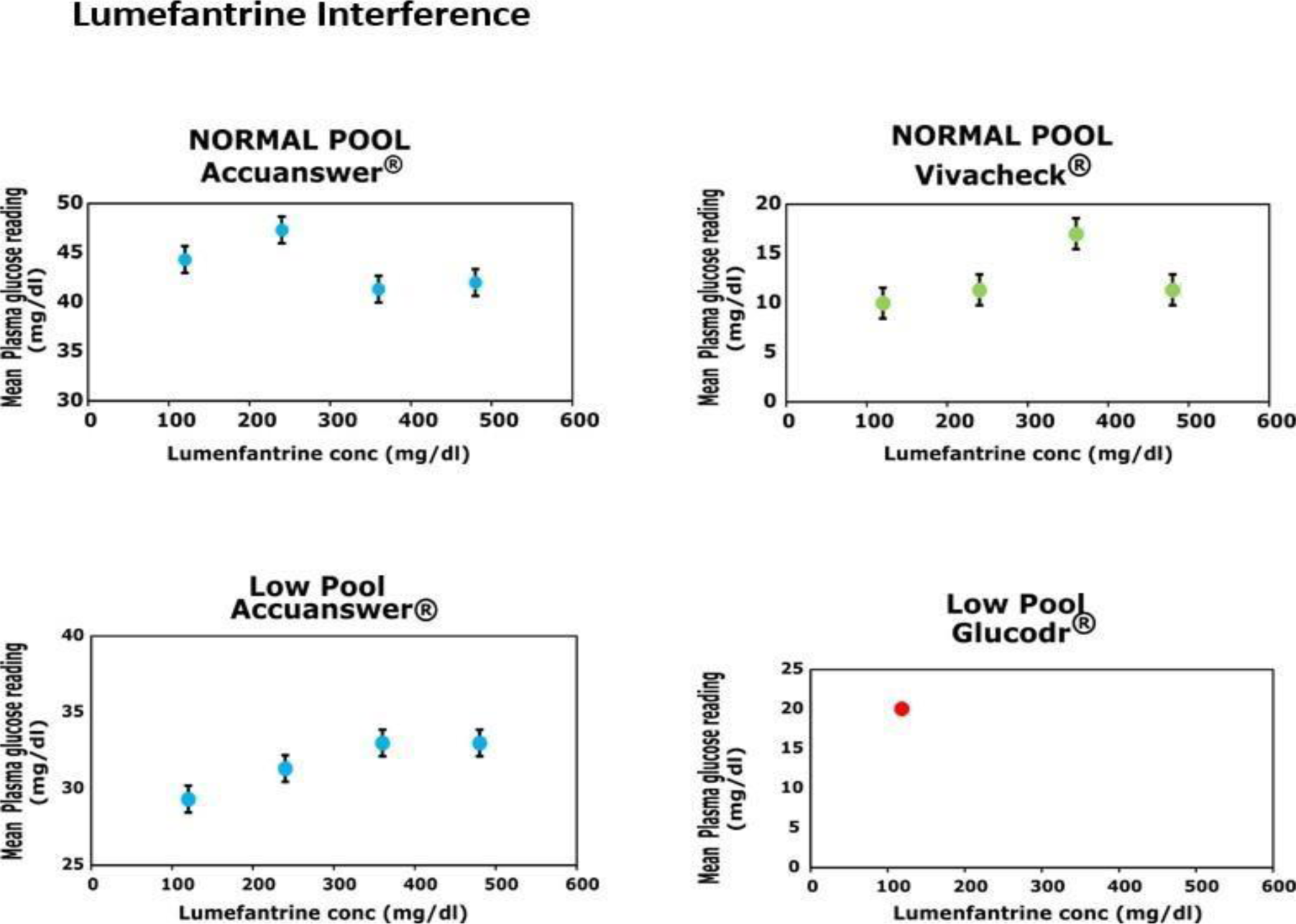
Influence of Lumefantrine in glucose level measurement in three models of glucometers –Vivacheck, Accu Answer and Glucodr – in normal and low pools.

These interferences by artemether and lumefantrine may provide false glucose readings in glucometer. The interference of glucose reading accuracy by some drugs have led to misdiagnoses according to previous studies [27]. Obviously, Accu-Answer^®^ glucometer consistently express artemether and lumefantrine interferences in glucose readings. This may justify the precision of the Accu -Answer^®^ glucometer which has a CoV less than 5%.

### 3.4 In silico analyses

Molecular docking is a computational biology method used to predict the binding affinities between a ligand-protein complexes and understand possible interactions between the ligands and the amino acid residues in the binding pockets of the target protein. In this study (Figure S4), we predicted for molecular interactions between artemether and lumefantrine, and the three glucometer enzymes in the binding pockets of glucose. In Table S1, artemether and lumefantrine showed low binding affinities to the three enzymes. However, lumefantrine showed higher binding affinities (−5.945, -5.043 and 3.726 kcal/mol) than artemether binding affinities (−5.469, -4.68 and -3.652 kcal/mol) in GOx (PDB: 1CF3), GDH-FAD (PDB: 4YNU) and GDH-PQQ (PDB: 1CQ1), respectively. The low binding affinities may be responsible for the mild interference of glucose reading accuracy that were observed in the three glucometers. Molecular docking analysis revealed that both artemether and lumefantrine interact with key amino acid residues of GOx such as TYR 80, 515, PHE 564, VAL 560, ILE 94, ALA 292, LEU 29, and MET 556 (Table S1). In addition, lumefantrine interacts with TYR 68, PHE 414, TRP 426, and HIS 516. Some of these amino acids are known to be potentially important for the binding of glucose (TYR 68, PHE 414, and TRP 426) as well as for the enzyme’s catalytic activity (HIS 516), indicating that the binding of lumefantrine to this binding pocket could be responsible for the high binding affinity compared to artemether.For GDH-FAD enzyme, we observed that artemether and lumefantrine has similar interactions amino acid residues such as TYR 53, 199, ALA 96, LEU 401, and MET 95. These amino acid residues in the binding poses are similar to the observed diffracted complex of glucose and GDH-FAD after X-ray diffraction [28]. Furthermore, the interactions of artemether and lumefantrine with GDH-PQQ indicate raised some concerns about the precision of the Accu -Answer glucometer. This was because there were no similar reports of the amino acid residues. However, a hallmark amino acid residue at the conserved activity site for glucose sensitivity in GOx and GDHis tyrosine 68 (Tyr68), which binds to the fourth carbon (C4) [26].

## 4. Conclusions

In summary, we validated the interference potentials of acetaminophen, ascorbic acid and maltose. We discovered that Vivachek^®^ glucometer may be a preferred glucometer for MID patients under A-L therapy because of the lack of interference in this glucometer. Nevertheless, Accu-Answer^®^ was observed to be the most precise glucometer in Nigeria. We found that artemether and lumefantrine could induce mild interference in Accu - Answer^®^ and Glucodr^®^ glucometers. The predicted binding poses of the molecular dockingmay inform protein engineering to optimize the specificity of the glucometer enzymes.

Distilled water was used to prepare the different solutions of the ACT drugs. However, the artemether and lumefantrine compounds did not dissolve completely in distilled water. Further studies are encouraged to investigate glucose readings with dissolved solutions of these drugs. Also, we encourage this study to be further investigated with MID patient in the clinic. In addition, the molecular docking was only a prediction and may not be theexact picture of the reality. Therefore, there is a need for an evaluation of the structural biology of the co-crystallized complexes (drugs and enzymes) through diffraction techniques.

## Supporting information

Supplemental list

## CRediT authorship contribution statement

**Ezinwanne Jane Ugochukwu**; Conceptualization, Methodology, Investigation, Writing original draft, Visualization. **Adaobi Uchenna Mosanya**; Conceptualization, Methodology, Investigation, Data curation, Formal analysis. **Victor Nnanna Nweke;** Conceptualization, Metodology, Data curation, review and edits **Marydith Ifeoma Chukwu**; Conceptualization, Methodology, Data curation, Formal analysis. **Chukwuemeka Michael Ubaka;** Conceptualization, Methodology, Supervision, Investigation, writing original draft, Visualization, review and edits **Mohammed Alawami**; Conceptualization, Methodology, Supervision, Investigation, Writing original draft, Visualization, review and edits.

## Acknowledgement

We thank the almighty God for his help throughout this program. We also thank the ReachSci society of the Cambridge University for the opportunity to partake in this ground breaking project. We also thank Emzor Pharmaceutical Industry and Neimeth Pharmaceutical PLC for providing us with some pure samples needed to carry out this research. Also, to our supervisor, Prof. Chukwuemeka Michael Ubaka for his encouragement, Prof. Ebere Onuigbo for helping us with some equipment needed for this research, Prof. A.A Attama for providing us with the pure samples of Artemether and Lumefantrine powders and finally to the team of University of Nigeria Nsukka for their relentless efforts in meeting up to deadlines and for being there to help in tasks even when it was difficult to show up. Funding by ReachSci society of the University of Cambridge is greatly acknowledged.

## References

[1] WHO (1999) WHO. Definition, diagnosis and classification of diabetes mellitus and itscomplications, part 1. Geneva: WHO; 1999 - Google Search. Available at: https://www.google.com/search?client=firefox-b-d&q=1.+WHO.+Definition%2C+diagnosis+and+classification+of+diabetes+mellitus+and+its+complications%2C+part+1.+Geneva%3A+WHO%3B+1999 (Accessed: 26December 2022).

[2] Uloko, A.E. et al. (2018) ‘Prevalence and Risk Factors for Diabetes Mellitus in Nigeria: ASystematic Review and Meta-Analysis’, Diabetes Therapy, 9(3), pp. 1307–1316. doi:10.1007/s13300-018-0441-1.

[3] IDF (2021) Diabetes Atlas: Diabetes in Africa. 10th ed. Brussels: International Diabetes Federation; 2021. - Google Search. Available at: https://www.google.com/search?q=3.%09International+Diabetes+Federation.+Diabetes+Atlas%3A+Diabetes+in+Africa.+10th+ed.+Brussels%3A+International+Diabetes+Federation%3B+2021.&client=firefox-b-d&ei=Tr2pY5CQCcaHkdUPw826gAI&ved=0ahUKEwiQ78amzJf8AhXGQ6QEHcOmDi (Accessed: 26 December 2022).

[4] Sharma, T. et al. (2014) ‘Poor adherence to treatment: A major challenge in diabetes’, Journal, Indian Academy of Clinical Medicine, 15(1), pp. 26–9.

[5] Osei-Yeboah, J. et al. (2018) ‘Physical Activity Patterns and Its Association with Glycaemic and Blood Pressure Control among People Living with Diabetes (PLWD) InThe Ho Municipality, Ghana’, Ethiop J Health Sci, 29(1), p. 819. doi:10.4314/ejhs.v29i1.3.

[6] Mikhael, E. et al. (2019) ‘Self-management knowledge and practice of type 2 diabetes mellitus patients in Baghdad, Iraq: A qualitative study’, Diabetes, Metabolic Syndrome and Obesity: Targets and Therapy, 12, pp. 1–17. doi:10.2147/DMSO.S183776.

[7] Chenoweth, J. et al. (2020) ‘Acetaminophen interference with Nova StatStrip^®^ GlucoseMeter: case report with bench top confirmation’, Clinical Toxicology, 58(11), pp. 1067–1070. doi:10.1080/15563650.2020.1732404.

[8] Calhoun, P. et al. (2018) ‘Resistance to Acetaminophen Interference in a Novel Continuous Glucose Monitoring System’, Journal of Diabetes Science and Technology, 12(2), pp. 393–396. doi:10.1177/1932296818755797.

[9] Zhang, Y. et al. (1994) ‘Elimination of the Acetaminophen Interference in an ImplantableGlucose Sensor’, Analytical Chemistry, 66(7), pp. 1183–1188. doi:10.1021/ac00079a038.

[10] Yoo, E. and Lee, S. (2010) ‘Glucose Biosensors: An Overview of Use in ClinicalPractice’, Sensors, 10, pp. 4558–4576. doi:10.3390/s100504558.

[11] World Health Organization (2021) ‘Global technical strategy for malaria 2016– 2030’, World Health Organization, pp. 1–35. Available at: https://apps.who.int/iris/bitstream/handle/10665/186671/9789243564999_spa.pdf?sequence=1.

[12] Ikekpeazu, E.J. et al. (2010) ‘Type-2 Diabetes Mellitus and Malaria Parasitaemia: Effect on Liver Function Tests’, Asian Journal of medical Sciences, 2(5), pp. 214–217.

[13] Fasanmade, O. and Dagogo-Jack, S. (2015) ‘Diabetes Care in Nigeria’, Annals ofGlobal Health, 81(6), pp. 821–829. doi:10.1016/j.aogh.2015.12.012.

[14] Atere, A. et al. (2020) ‘Correlation between Anemia and Malaria Infection Severity inPatients with Type 2 Diabetes Mellitus in Nigeria’, Althea Medical Journal, 7(4), pp. 170–175. doi:10.15850/amj.v7n4.2085.

[15] Udoh, B. et al. (2020) ‘Asymptomatic falciparum malaria and its effects on type 2 diabetes mellitus patients in Lagos, Nigeria’, Saudi Journal of Medicine and MedicalSciences, 8(1), p. 32. doi:10.4103/sjmms.sjmms_178_18.

[16] Ibrahim, A. et al. (2022) ‘Socio-Demographic Profile, Asymptomatic Malaria Parasitaemia and Glycemic Control among Midled-Aged and Elderly Type 2 Diabetes Mellitus Patients in Rural Southwestern Nigeria: A Cross Sectional Study’, African Journal of Diabetes Medicine, 30(3), pp. 1–7. doi:10.54931/2053-4787.30-3-1.

[17] Cho, J. et al. (2016) ‘Influence of Vitamin C and Maltose on the Accuracy of ThreeModels of Glucose Meters’, Annals of Laboratory Medicine, 36, pp. 271–274.

[18] Olsson, M. et al. (2011) ‘PROPKA3: Consistent treatment of internal and surface residues in empirical p K a predictions’, Journal of Chemical Theory and Computation,7(2), pp. 525–537. doi:10.1021/ct100578z

[19] Wong, C., Wong, K. and Chen, X. (2008) ‘Glucose oxidase: Natural occurrence, function, properties and industrial applications’, Applied Microbiology and Biotechnology, 78(6), pp. 927–938. doi:10.1007/s00253-008-1407-4

[20] Vanavanan, S. et al. (2010) ‘Performance of a new interference-resistant glucose meter’, Clinical Biochemistry, 43(1–2), pp. 186–192. doi:10.1016/j.clinbiochem.2009.09.010.

[21] Tang, Z. et al. (2000) ‘Effects of pH on glucose measurements with handheld glucose meters and a portable glucose analyzer for point-of-care testing’, Archives of Pathologyand Laboratory Medicine, 124(4), pp. 577–582. doi:10.5858/2000-124-0577-eopogm.

[22] Sartor, Z., Kesey, J. and Dissanaike, S. (2015) ‘The effects of intravenous vitamin C on point-of-care glucose monitoring’, Journal of Burn Care and Research, 36(1), pp. 50–56.doi:10.1097/BCR.0000000000000142.

[23] Cartier, L.J. et al. (1998) ‘Toxic levels of acetaminophen produce a major positive interference on Glucometer Elite and Accu -chek Advantage glucose meters.’, Clinicalchemistry. England, pp. 893–894.

[24] Maahs, D.M. et al. (2015) ‘Effect of Acetaminophen on CGM Glucose in an OutpatientSetting’, 38. doi:10.2337/dc15-1096.

[25] Janssen, W. et al. (1998) ‘Positive interference of icodextrin metabolites in someenzymatic glucose methods [4]’, Clinical Chemistry, 44(11), pp. 2379–2380. doi:10.1093/CLINCHEM/44.11.2379.

[26] Ferri, S., Kojima, K. and Sode, K. (2011) ‘Review of glucose oxidases and glucose dehydrogenases: A bird’s eye view of glucose sensing enzymes’, Journal of DiabetesScience and Technology, 5(5), pp. 1068–1076. doi:10.1177/193229681100500507.

[27] Bauer, J. et al. (2022) ‘Glucose Oxidase, an Enzyme “Ferrari”: Its Structure, Function,Production and Properties in the Light of Various Industrial and Biotechnological Applications’, Biomolecules, 12(3), p. 472. doi:10.3390/BIOM12030472.

[28] Yoshida, H. et al. (2015) ‘Structural analysis of fungus-derived FAD glucosedehydrogenase’, Scientific Reports, 5(1), pp. 1–13. doi:10.1038/srep1349.

