## Supplemental list for "Study of Glucometer Precision and Antimalarial Therapy Interference on Glucose Readings in Nigeria"

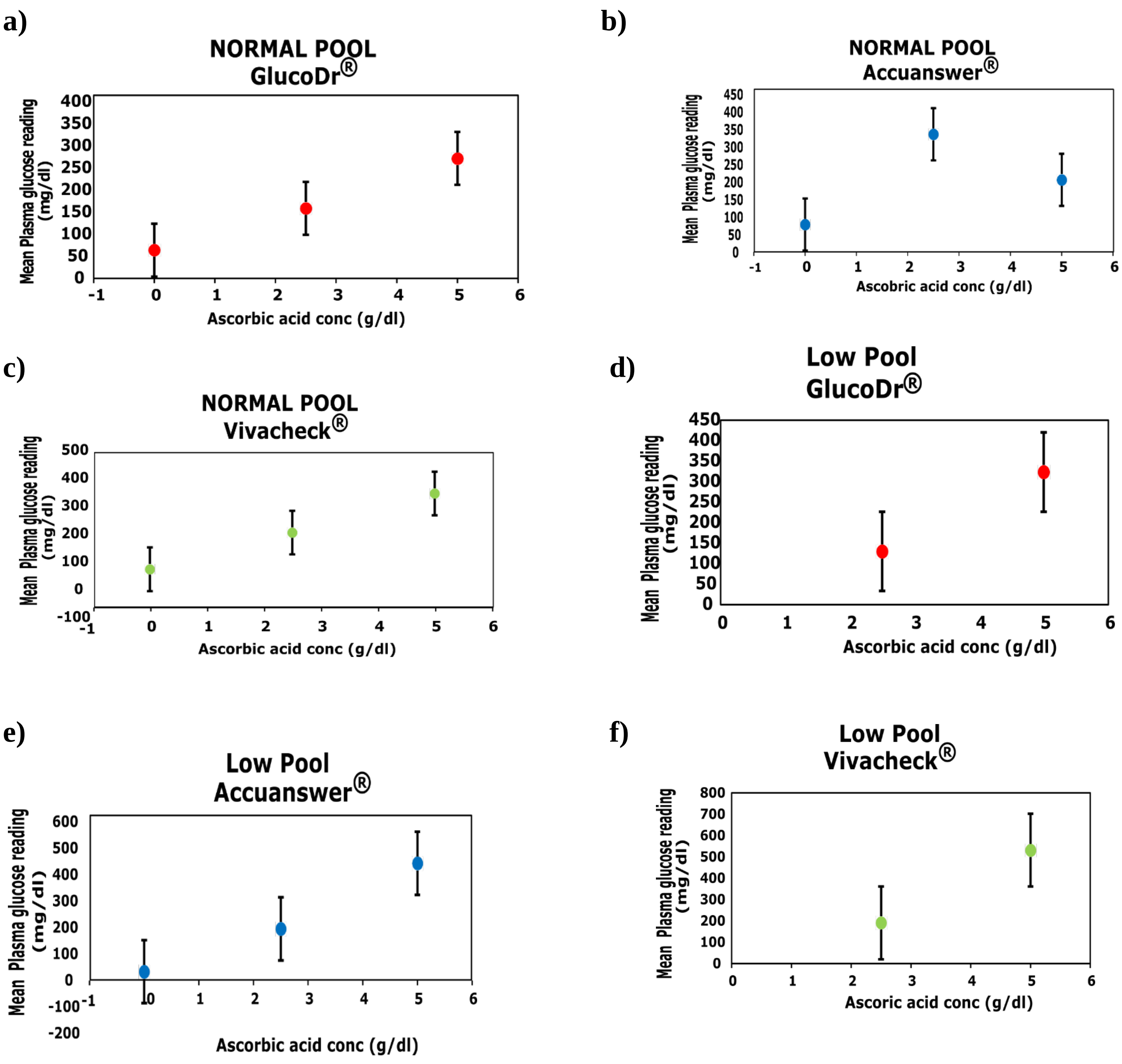


**Figure S1: a)** Influence of Ascorbic acid on mean plasma glucose reading in GlucoDr® glucometers in normal pool**. b)** Influence of Ascorbic acid on mean plasma glucose reading in Accuanswer® glucometers in normal pool. **c)** Influence of Ascorbic acid on mean plasma glucose reading in Vivacheck® glucometers in normal pool. **d)** Influence of Ascorbic acid on mean plasma glucose reading in GlucoDr® glucometers in low pool. **e)** Influence of Ascorbic acid on mean plasma glucose reading in Accuanswer® glucometers in low pool**. f)** Influence of Ascorbic acid on mean plasma glucose reading in Vivacheck® glucometers in low pool.

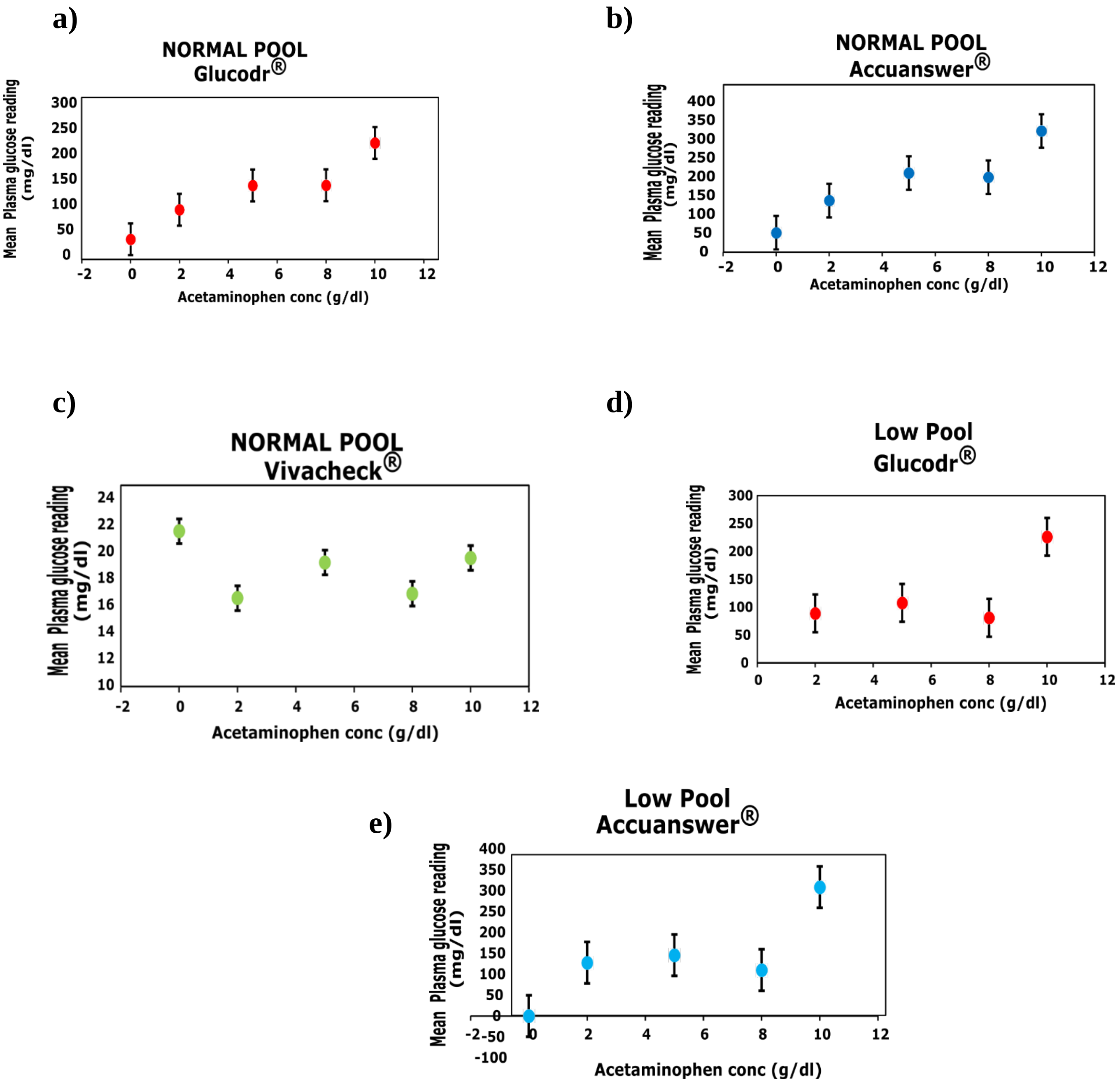


**Figure S2: a)** Influence of Acetaminophen on mean plasma glucose reading in GlucoDr® glucometers in normal pool**. b)** Influence of Acetaminophen on mean plasma glucose reading in Accuanswer® glucometers in normal pool. **c)** Influence of Acetaminophen on mean plasma glucose reading in Vivacheck® glucometers in normal pool. **d)** Influence of Acetaminophen on mean plasma glucose reading in GlucoDr® glucometers in low pool. **e)** Influence of Acetaminophen on mean plasma glucose reading in Accuanswer® glucometers in low pool**.**

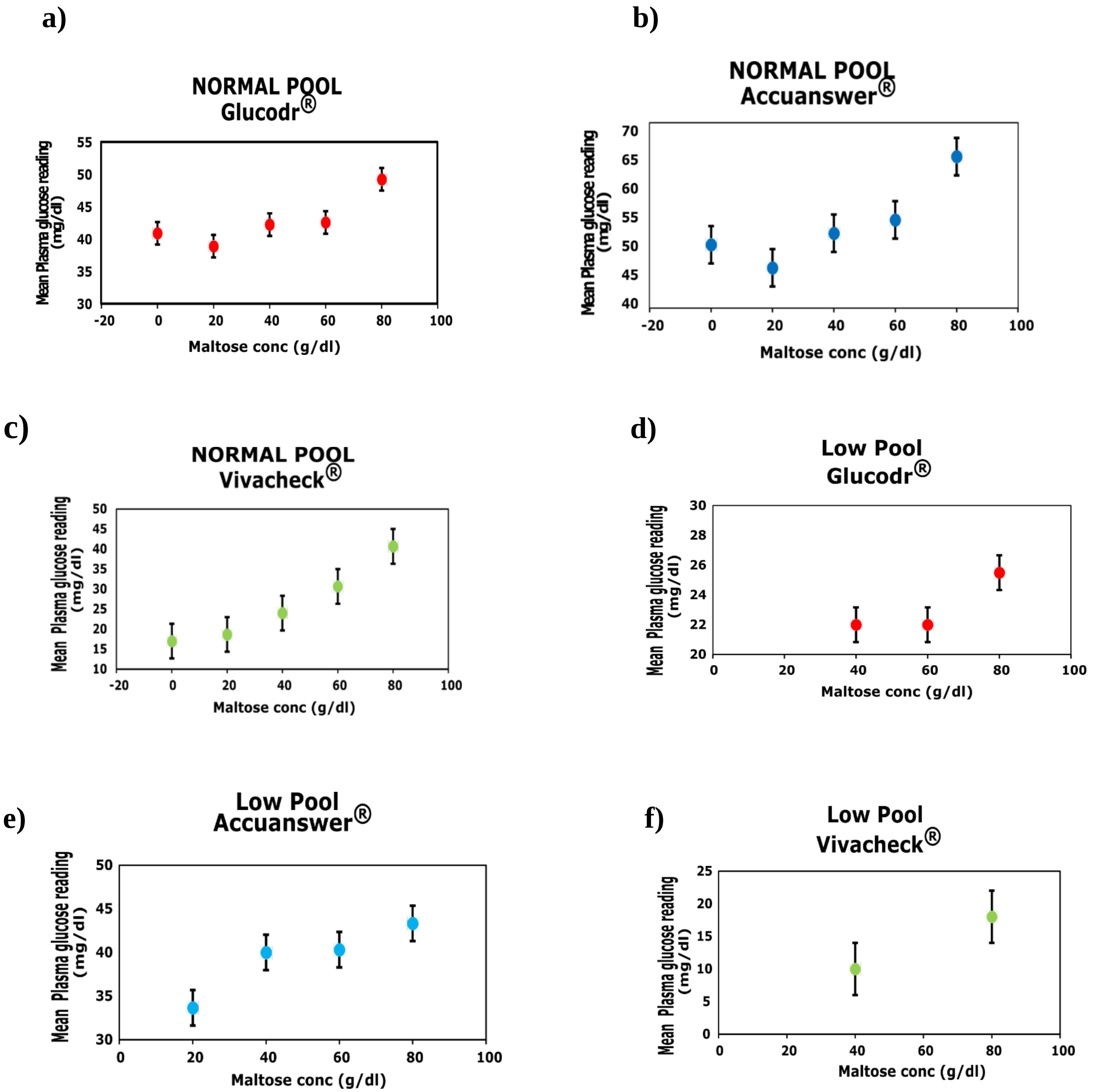


**Figure S3: a)** Influence of Maltose on mean plasma glucose reading in GlucoDr® glucometers in normal pool**. b)** Influence of Maltose on mean plasma glucose reading in Accuanswer® glucometers in normal pool. **c)** Influence of Maltose on mean plasma glucose reading in Vivacheck® glucometers in normal pool. **d)** Influence of Maltose on mean plasma glucose reading in GlucoDr® glucometers in low pool. **e)** Influence of Maltose on mean plasma glucose reading in Accuanswer® glucometers in low pool**. f)** Influence of Maltose on mean plasma glucose reading in Vivacheck® glucometers in low pool.

a)

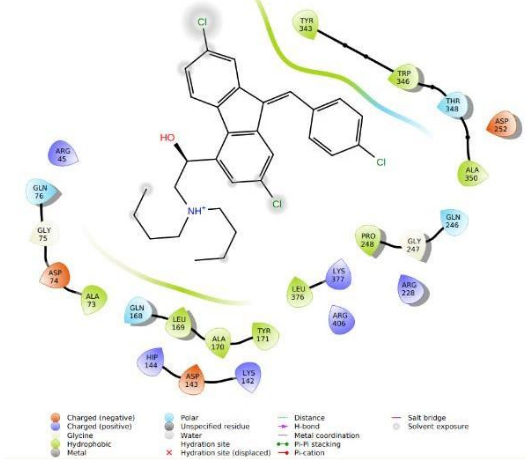
Cl
HO
NH+
TYR 343
TRP 346
THR 348
ASP 252
ALA 350
ARG 45
GLN 76
GLY 75
ASP 74
ALA 73
GLN 246
GLY 247
PRO 248
LYS 377
LEU 376
ARG 228
ARG 406
GLN 168
LEU 169
ALA 170
TYR 171
HIP 144
ASP 143
LYS 142
Charged (negative)
Charged (positive)
Glycine
Hydrophobic
Metal
Polar
Unspecified residue
Water
Hydration site
Hydration site (displaced)
Distance
H-bond
Metal coordination
Pi-Pi stacking
Pi-cation
Salt bridge
Solvent exposure

b)

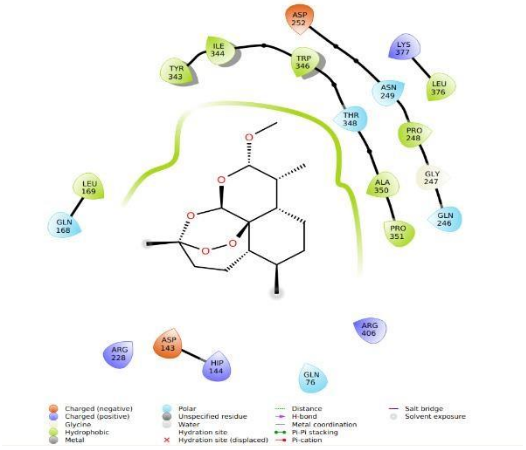
ASP 252
TYR 343
ILE 344
TRP 346
LYS 377
LEU 376
ASN 249
THR 348
PRO 248
GLY 247
LEU 169
GLN 168
ALA 350
GLN 246
PRO 351
ARG 406
ARG 228
ASP 143
HIP 144
GLN 76
Charged (negative)
Charged (positive)
Glycine
Hydrophobic
Metal
Polar
Unspecified residue
Water
Hydration site
Hydration site (displaced)
Distance
H-bond
Metal coordination
Pi-Pi stacking
Pi-cation
Salt bridge
Solvent exposure

c)

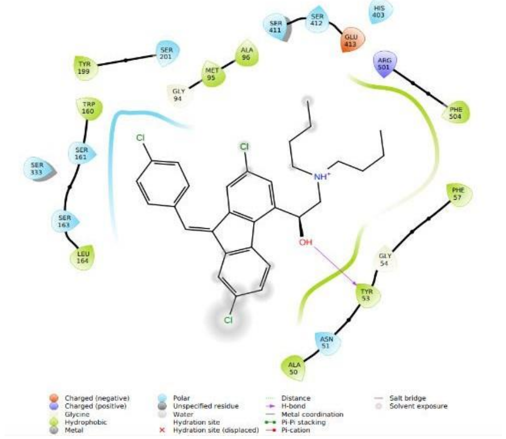
SER 411
SER 412
HIS 403
GLU 413
TYR 199
SER 201
MET 95
ALA 96
ARG 501
GLY 94
TRP 160
PHE 504
SER 161
SER 333
SER 163
LEU 164
PHE 57
GLY 54
TYR 53
ASN 51
ALA 50
Cl
NH+
OH
Charged (negative)
Charged (positive)
Glycine
Hydrophobic
Metal
Polar
Unspecified residue
Water
Hydration site
Hydration site (displaced)
Distance
H-bond
Metal coordination
Pi-Pi stacking
Pi-cation
Salt bridge
Solvent exposure

d)

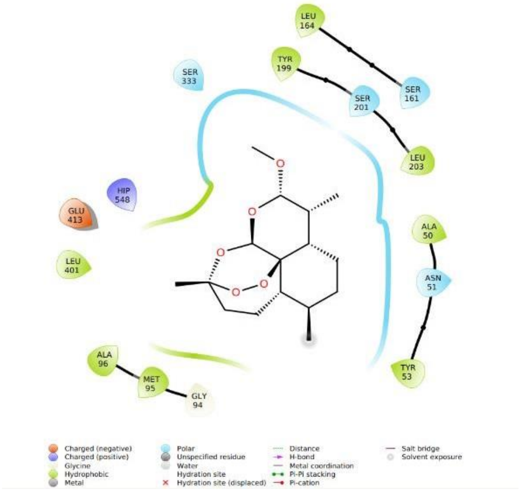
LEU 164
TYR 199
SER 333
SER 161
SER 201
LEU 203
HIP 548
GLU 413
ALA 50
LEU 401
ASN 51
TYR 53
ALA 96
MET 95
GLY 94
Charged (negative)
Charged (positive)
Glycine
Hydrophobic
Metal
Polar
Unspecified residue
Water
Hydration site
Hydration site (displaced)
Distance
H-bond
Metal coordination
Pi-Pi stacking
Pi-cation
Salt bridge
Solvent exposure

e)

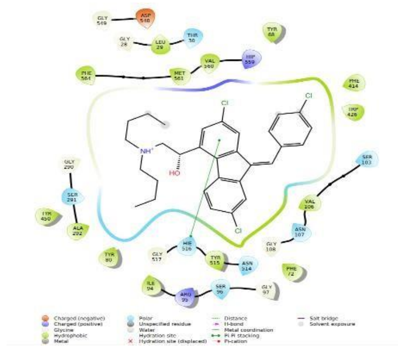

f)

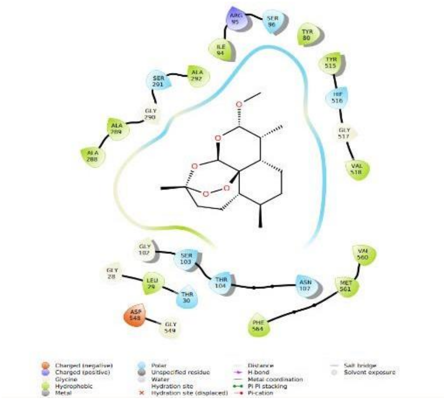

**Figure S4: a)** 2D representations of molecular docking interactions of artemether with Glucose dehydrogenase FAD. **b)** 2D representations of molecular docking interactions of lumenfantrine with Glucose dehydrogenase FAD. **c)** 2D representations of molecular docking interactions of artemether with Glucose dehydrogenase PQQ. **d)** 2D representations of molecular docking interactions of lumenfantrine with Glucose dehydrogenase PQQ. **e)** 2D representations of molecular docking interactions of artemether with Glucose oxidase. **f)** 2D representations of molecular docking interactions of lumefantrine with Glucose oxidase.

| Target | Compound | Docking Score | Hydrogen Bond Interaction | Hydrophobic Interaction | Pi-Pi Stacking Interaction |
|---|---|---|---|---|---|
| **1CF3** | Lumefantrine | -5.945 | | TYR 68, 80, 450, 515, PHE 72, 414, 564, VAL 106, 560, ALA 292, ILE 94, LEU 29, MET 561, TRP 426 | HIE 516 |
| | Artemether | -5.469 | | TYR 80, 515, PHE 564, VAL 518, 560, ILE 94, ALA 288, 289, 292, LEU 29, MET 561 | |
| **4YNU** | Lumefantrine | -5.043 | TYR 53 | TYR 53, 199, ALA 50, 96, PHE 57, 504, TRP 160, LEU 164, MET 95 | |
| | Artemether | -4.68 | | TYR 53, 199, ALA 50, 96, LEU 164, 203, 401, MET 95 | |
| **1CQ1** | Lumefantrine | -3.726 | | TYR 171, 343, ALA 350, TRP 346, PRO 248, LEU 376, 169 | |
| | Artemether | -3.652 | | TYR 343, ALA 350, TRP 346, PRO 248, 351, LEU 376, 169, ILE 344 | |

**Table S1:** Docking scores of the biological interactions of artemether and lumenfantrine with GOx (PDB: 1CF3), GDH-FAD (PDB: 4YNU) and GDH-PQQ (PDB: 1CQ1).
